# Cytosolic proteins can exploit membrane localization to trigger functional assembly

**DOI:** 10.1101/164152

**Authors:** Osman N. Yogurtcu, Margaret E. Johnson

## Abstract

Cell division, endocytosis, and viral budding would not function without the localization and assembly of protein complexes on membranes. What is poorly appreciated, however, is that by localizing to membranes, proteins search in a reduced space that effectively drives up concentration. Here we derive an accurate and practical analytical theory to quantify the significance of this dimensionality reduction in regulating protein assembly on membranes. We define a simple metric, an effective equilibrium constant, that allows for quantitative comparison of protein-protein interactions with and without membrane present. To test the importance of membrane localization for driving protein assembly, we collected the protein-protein and protein-lipid affinities, protein and lipid concentrations, and volume-to-surface-area ratios for 46 interactions between 37 membrane-targeting proteins in human and yeast cells. We find that many of the protein-protein interactions between pairs of proteins involved in clathrin-mediated endocytosis in human and yeast cells can experience enormous increases in effective protein-protein affinity (10-1000 fold) due to membrane localization. Localization of binding partners thus triggers robust protein complexation, suggesting that it can play an important role in controlling the timing of endocytic protein coat formation. Our analysis shows that several other proteins involved in membrane remodeling at various organelles have similar potential to exploit localization. The theory highlights the master role of phosphoinositide lipid concentration, the volume-to-surface-area ratio, and the ratio of 3D to 2D equilibrium constants in triggering (or preventing) constitutive assembly on membranes. Our simple model provides a novel quantitative framework for interpreting or designing *in vitro* experiments of protein complexation influenced by membrane binding.

## INTRODUCTION

When clathrin, the essential cytosolic protein of clathrin-mediated endocytosis (CME), self-assembles into multi-protein cages, the same protein-protein contacts are used regardless of whether clathrin is in solution or on the membrane (1-3). So why is more binding (2) observed on the membrane? The reason is dimensionality reduction: proteins on the membrane search a smaller space and this increases their relative concentration; higher concentration of proteins promotes binding simply due to LeChatelier’s principle. The question we address is, how significant a role can this dimensionality reduction play for driving protein-protein interactions between cytosolic proteins *in vitro*? Understanding this role can help interpret mechanisms of assembly *in vivo*. Despite the wide-ranging cellular processes such as cell division and viral budding that could exploit this phenomenon, it has so far lacked a predictive theoretical framework. Hence while the concept that membrane localization can enhance binding may be familiar or intuitive, we here make that concept quantitative for soluble binding partners. In contrast, theory for understanding reduction of dimensionality in chemoreception and receptor mediated signaling (where it can also be functionally significant (4)) has been studied for decades (5, 6). Membrane localization can accelerate a ligand’s search for membrane bound targets (5-9) and increase activation of intracellular receptors, influencing downstream response (8-10). However, in these cases, a soluble protein always targets a membrane bound receptor. Here we capture the dynamic cases where both binding partners are soluble and target lipids present in limited concentrations, as occurs, for example, in CME. Our theory determines how binding enhancement depends on protein and lipid concentrations, protein-protein and protein-lipid affinities, the volume-to-surface area ratio, and the change in binding affinities from 3D to 2D. Quantifying this behavior is critical to understanding assembly on surfaces because 2D localization can strengthen binding reactions regardless of whether additional factors, such as curvature generation (11), membrane microdomains (12, 13), or conformational switches (1), also influence binding.

We show here that membrane localization offers a powerful way of controlling protein concentrations and therefore of regulating the timing of multi-protein assembly. The analytical theory we present describes a relatively simple model at equilibrium where a pair of soluble binding partners can form complexes in solution and also can both bind and continue to form complexes on the surface of a membrane (Figure 1). Thus it is useful as a tool to quantify protein-protein interactions that, while physiologically relevant, are being studied *in vitro.* Without accounting for the complex array of factors present *in vivo,* such as variability in membrane composition, competition for protein and lipid binding from diverse proteins, spatial distributions of proteins or lipids, and non-equilibrium dynamics, we can only speculate about the behavior in the cell. However, the theory provides a novel and valuable metric for interpreting how important membrane localization can be given the concentrations and binding properties of component proteins, and it isolates the role of membrane localization from other factors. Since even most *in vitro* experiments contain more components and complexity than is captured in our simple model, we discuss how it can still be used as a quantitative guide for estimating how membrane heterogeneity, competition for binding, and mutations would influence the parameters of the model (volume, surface area, binding affinities and concentrations) and thus the proteins’ subsequent response to localization. We specifically address in our results how lipids such as PI(4,5)P_2_ can be targeted by many proteins at any time (14, 15), how some membrane binding domains such as BAR domains bind membranes with widely varying lipid composition and in a curvature dependent manner (16, 17), and how mutations and multiple protein binding partners would alter protein complex formation. Despite the limitations of applying an equilibrium theory to understand complexation that ultimately occurs in the nonequilibrium cell, we believe the theory represents a well--defined starting point from which to probe more complex systems, just as using *in vitro* studies provide a useful guide for interpreting behavior in the cell. It is also a reference point for studying the time-dependence of assembly through computer simulation, as we do here, and a starting point from which to build further complexity into the model.

**Figure 1:**
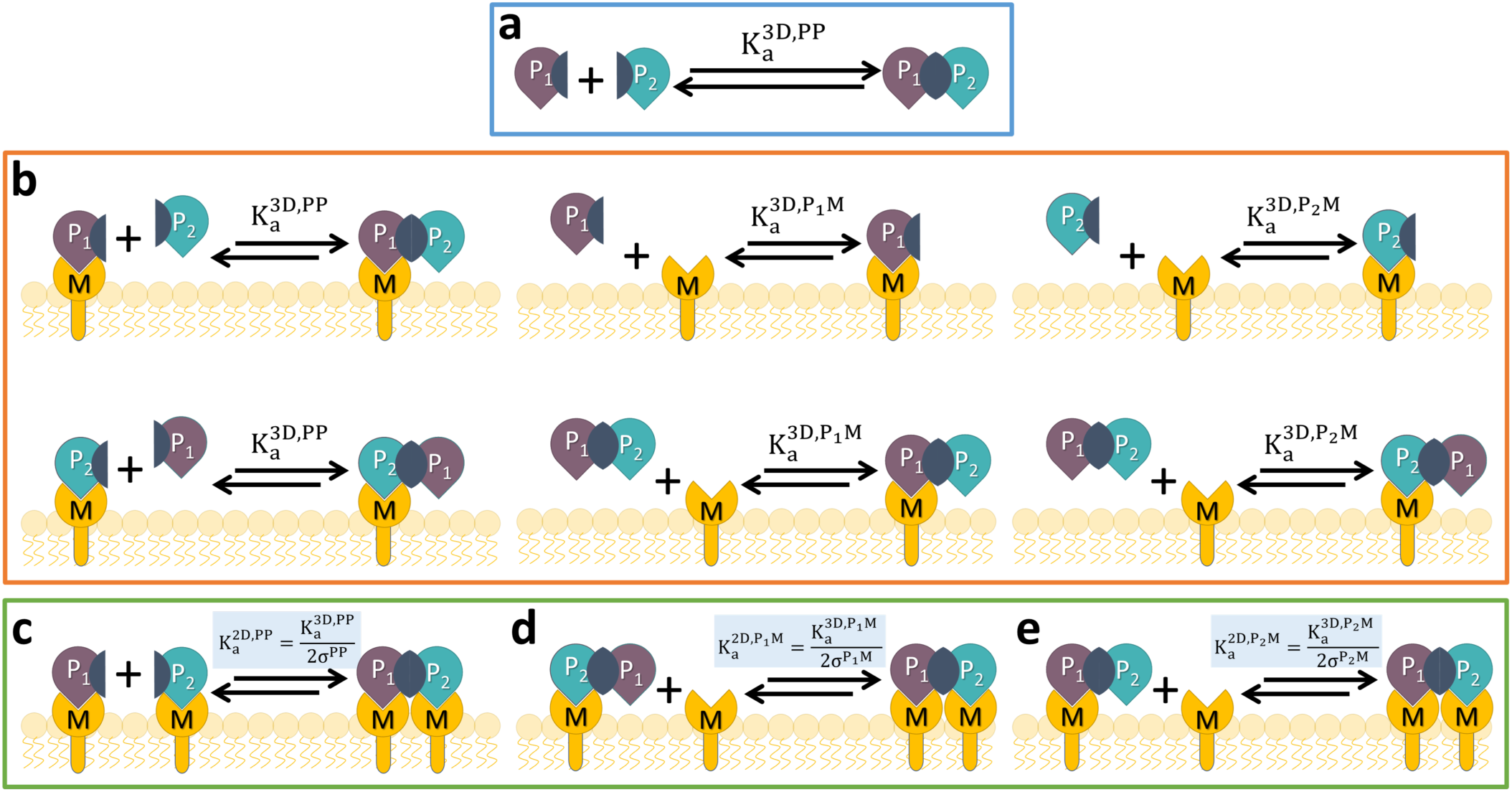
Quantifying how protein binding partners in solution can increase complex formation by binding to lipids on membrane surfaces. We show all the possible binding interactions that can occur between a pair of proteins that bind each other (P_1_+P_2_⇌P_1_P_2_) and also bind specific lipids (M). a) Solution (3D) binding. b) Interactions in solution (3D) that pull proteins to the membrane surface through protein or lipid binding. In (c-e) the binding interactions are in 2D (species concentration in Area-1) and can exploit the lower search space. Conversion from 3D to 2D equilibrium constant is defined by the variable σ, where only s=sPP appears in Eq. 3. To solve for all species in consistent units (i.e. Volume^−1^), K_a_^2D^ values must be multiplied by V/A. The size of solution volume V vs. membrane surface area A is thus a critical parameter controlling binding enhancement. The membrane surface can be the plasma membrane, for example, but also liposomes suspended in solution. d,e) Proteins localized at the surface will also bind lipids in a 2D search. There are over 100 functionally diverse peripheral membrane proteins in yeast alone (14) whose binding interactions with one another could strengthen substantially via binding to membranes. We simulated this model for a comprehensive range of conditions using mainly systems of ordinary differential equations (ODE), but also single-particle reaction-diffusion (RD) simulations (19, 20) (Methods).

We apply the theory here to characterizing, within a quantitative framework, the role of membrane localization for enhancing 55 binding interactions involving 33 distinct protein pair interaction sets (Table S1). Through simulation, we also move beyond the model illustrated in Figure 1 of only pairs of soluble binding partners to show how complexation involving non-membrane binding scaffold proteins such as clathrin, or how formation of higher-order oligomers, which is functionally important for driving membrane remodeling (11, 17, 18), can also be regulated by membrane localization (13 additional interaction sets in Table S2). Our theory only applies to the pair interactions illustrated in Figure 1. We include 22 proteins involved in CME in both human and yeast cells, as well as 15 proteins involved in lipid regulation, vesicle formation on endosomes, budding, and morphogenesis in yeast cells (Table 1). We collected concentration and cellular geometry data based on *in vivo* values to better connect to physiologic regimes. Although our theory represents an approximate solution to the full model shown in Figure 1, we show through extensive simulations using both systems of ordinary differential equations and single-particle reaction-diffusion (19, 20) that it is highly accurate. Through simulation, we additionally find that membrane localization alters the timescales of protein-rotein assembly, but that the result is not dominated by changes in protein diffusion between solution and the membrane. Rather, for physiologic binding strengths, the rate-limiting step is the speed of binding to the membrane surface from solution. Finally, a practical application of our simple formula is that it can be used to experimentally fit protein-protein binding affinities on surfaces (K_a_^2D^), which are rarely measured (21, 22). The advantage of the formula is that it applies to *in vitro* experiments where a pair of proteins can *reversibly* bind to the membrane, thus avoiding the need to restrict proteins to the surface.

**Table 1.** Proteins studied, copy numbers, protein-lipid interactions (PLI) and affinities (K_d_^PM^)

| | Protein | Species | Copy Number<sup>d</sup> | $K_d^{PM}$ (μM) | Literature Refs |
| --- | --- | --- | --- | --- | --- |
| 1 | OSH2 | Yeast | 850 | 1-1.5 | PLI: PMID:11238399. Affinity, measured: same ref. |
| 2 | SWH1 | Yeast | 505 | 3.5-6.2 | PLI: PMID:21119626. Affinity, measured: same ref. |
| 3 | KES1 | Yeast | 21166 | 0.055-2.2 | PLI: PMID:22162133,11916983. Affinity, measured: same. |
| 4 | VPS17 | Yeast | 1077 | >100 | PLI: PMID:11557775. Affinity, measured: same ref. |
| 5 | SNX4 | Yeast | 1483 | >100 | PLI: PMID:11557775. Affinity, measured: same ref. |
| 6 | SNX41 | Yeast | 367 | >100 | PLI: PMID:11557775. Affinity, measured: same ref. |
| 7 | VPS5 | Yeast | 1326 | >100 | PLI: PMID:11557775. Affinity, measured: same ref. |
| 8 | ATG20 | Yeast | 519 | >100 | PLI: PMID:11557775. Affinity, measured: same ref. |
| 9 | BOI2 | Yeast | 567 | 6.6-19.5 | PLI: PMID:15023338. Affinity, measured: same ref. |
| 10 | CLA4 | Yeast | 397 | 20.2-100 | PLI: PMID:15023338. Affinity, measured: same ref. |
| 11 | SKM1 | Yeast | 16 | 3.9-6.4 | PLI: PMID:15023338. Affinity, measured: same ref. |
| 12 | BEM1 | Yeast | 1037 | >100 | PLI: PMID:11557775. Affinity, measured: same ref. |
| 13 | BOI1 | Yeast | 399 | 20 | PLI: PMID:15023338. Affinity, measured: same ref. |
| 14 | VAM7 | Yeast | 210 | 2-3 | PLI: PMID:11557775. Affinity, measured: same ref. |
| 15 | SNX3 | Yeast | 5092 | 2-3 | PLI: PMID:11557775. Affinity, measured: same ref. |
| 16 | SLA2 | Yeast | 3904 | 0.27-3.4 | PLI: PMID:15574875. Affinity, homology (AP180): PMID:12740367. |
| 17 | SYP1 | Yeast | 2467 | Used 0.1, 10, 100 | PLI: PMID:19713939,1321812. Affinity, not known, estimated. |
| 18 | ENT1 | Yeast | 1750 | 0.08 | PLI: PMID:22193158,10449404. Affinity, homology (EPN1): PMID:17825837. |
| 19 | ENT2 | Yeast | 1325 | 0.08 | PLI: PMID:22193158,10449404. Affinity, homology (EPN1): PMID:17825837. |
| 20 | YAP1802 | Yeast | 264 | 0.27-3.4 | PLI: PMID:22193158,21119626. Affinity, homology (AP180): PMID:12740367. |
|  | CHC1 (Not Studied) | Yeast | 19278<sup>a</sup> | No binding | - |
| 21 | SLA1 | Yeast | 2964 | No binding | - |
| 22 | EDE1 | Yeast | 5964 | No binding | - |
| 23 | FCHO1 | Human | 3706 | Used 0.1, 10, 100 | PLI: PMID:22484487. Affinity, not known, estimated. |
| 24 | AP-2 | Human | 244537<sup>b</sup> | 2.86 (0.072) | PLI: PMID:15916959. Affinity, measured: same ref. |
| 25 | EPN1 | Human | 570949 | 0.08 | PLI: PMID:17825837. Affinity, measured: same ref. |
| 26 | PICALM | Human | 358673 | 2.7-3.4 | PLI: PMID:25090048. Affinity, homology (AP180): PMID:12740367. |
| 27 | DAB2 | Human | 1078162 | 0.08 | PLI: PMID:12234931. Affinity, homology (EPN1): PMID:17825837. |
| 28 | FCHO2 | Human | 36302 | Used 0.1, 10, 100 | PLI: PMID:20448150. Affinity, not known, estimated. |
| 29 | SNAP91/AP180 | Human | 21716<sup>c</sup> | 0.27-3.4 | PLI: PMID:12740367. Affinity, measured: same ref. |
| 30 | LDLRAP1/ARH | Human | 1048 | 0.08 | PLI: PMID:12234931. Affinity, homology (EPN1): PMID:17825837. |
| 31 | HIP1 | Human | 13771 | 0.27-3.4 | PLI: PMID:14732715. Affinity, homology (AP180): PMID:12740367. |
| 32 | HIP1R | Human | 24161 | 0.27-3.4 | PLI: PMID:14732715. Affinity, homology (AP180): PMID:12740367. |
| 33 | AMPH | Human | 89536 | 0.1 | PLI: PMID:22888025. Affinity, measured: same ref. |
| 34 | SH3GL2/Endophilin | Human | 55621 | 0.03 | PLI: PMID:22888025. Affinity, measured: same ref. |
| 35 | EPS15 | Human | 91354 | No binding | - |
| 36 | ITSN1 | Human | 20184 | No binding | - |
| 37 | CLTC | Human | 1495814<sup>a</sup> | No binding | - |

a) To simulate clathrin trimers, the heavy chain copy numbers reported here are divided by 3.
- b) Considering AP2A1 gene. - c) Based on ppm value from PMID: 24920484. We scaled this value by the number of AP-2s from PMID: 26496610 to obtain the predicted number of AP180s in the cell. - d) Copy number for yeast from Ref. (45) and humans Ref: (46)

## MODEL and THEORY

In our primary model, we consider two proteins P_1_ and P_2_, that bind in solution with equilibrium constant 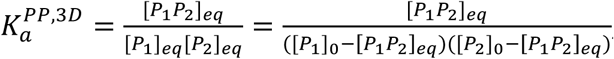, where total concentrations [*P*_1_]= [*P*_1_*P*_2_] + [*P*_1_]of the proteins are and same for [P_2_]_0_. If these proteins can also reversibly bind to membranes via targeting a specific lipid *M*, and continue to bind one another, their binding equilibrium will shift as a total of nine distinct species can form (Figure 1, Methods). The bound protein-protein complexes can either be in solution or on the membrane, [*P*_1_*P*_2_] ^sol + mem^ =[ *P*_1_*P*_2_] ^sol^ +[ *P*_1_*P*_2_] ^mem^ = [*P*_1_*P*_2_]+[*P*_1_*P*_2_*M*]+[*MP*_1_*P*_2_]+[*MP*_1_*P*_2_*M*], and unbound species are similarly defined =[P_1_]^sol^+[P_1_]^mem^=+[P_1_] ^sol^ +[MP_1_], and =[P_2_] +[P_2_M],where *M* indicates a copy of a target membrane lipid bound to P_1_ or P_2_. The model thus assumes each protein binds membrane via targeting a single copy of a specific lipid type. Proteins on the membrane must be able to move to bind one another, which is consistent with experimental observations (23) even of RNA-protein complexes (>6000kDa) that are anchored via multiple lipid binding sites along with myristoyl groups (24). Each of the nine distinct species will be constrained to preserve detailed balance at equilibrium, as defined by the 10 pairwise binding interactions of Figure 1 (see Methods), and the total concentrations of proteins is fixed at the same values as above, but now [*P*_1_]_0_=[*P*_1_*P*_2_] ^sol+mem^+[*P*_1_] ^sol+mem^ and the same for. Similarly,+[M]_0_+[M]+ [*P*_1_*P*_2_*M*]+ [*MP*_1_*P*_2_] 2[*MP*_1_*P*_2_*M*]. We note that species on the membrane have concentrations normally of μm^−2^, matching the units of equilibrium constants in 2D (K_a_^2D^)^−1^. All species copy numbers, whether on or off the membrane, however, can be solved for in volume units when the appropriate Solution volume/Membrane surface Area (V/A) conversion factor scales the 2D binding constants, so we always report volume units for concentrations. To quantify the change in bound protein-protein complexes as a function of membrane localization we will define an effective equilibrium constant

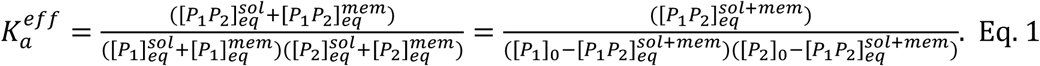

This is not a true equilibrium constant, as both bound and unbound states as defined above contain several species that do not all stepwise interconvert with one another. However, from K_a_^eff^ and initial protein concentrations [*P*_1_]_0_ and [*P*_2_]_0_, one can immediately solve for bound complex concentration using Eq. 1. If proteins cannot bind to the membrane, the value of K_a_^eff^ will revert to the solution bound value, K_a_^PP^, and thus the ratio of K_a_^eff^ / K_a_^PP^ determines the extent to which membrane localization either enhances or diminishes protein-protein complex formation. As we discuss further below, in the extreme limits where all proteins are either in solution or on the membrane, K_a_^eff^ reduces to a true equilibrium constant. The strength of K_a_^eff^ is that it also quantitatively describes all the conditions in between these limits. Thus, our K_a_^eff^ definition offers a valuable metric for quantifying the equilibrium of the model in Figure 1, which must otherwise be defined by multiple quantities.

We derive below an exact expression for K_a_^eff^ based on the 10 individual equilibrium relations for each reversible binding reaction (Figure 1, Methods). The value of K_a_^eff^ for any protein pair will depend on volume V, surface area A, total protein [P_1_]_0_, [P_2_]_0_, and lipid concentrations [M]_0_, and all six true equilibrium constants between protein and lipid interactions in 3D 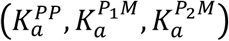 and in 2D 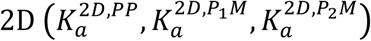 Importantly, the K_a_^2D^ values (with units μm2/mol, e.g.) are different from the corresponding 3D values, but they are related through K_a_^2D^=K_a_^3D^/(2σ). The variable σ, with units of length, is a thermodynamic property of each binding pair that captures largely entropic changes to binding as a result of surface restriction (22), as well as changes in the standard state units. It thus represents an independent variable that is specific to each protein pair studied. It is possible that even if concentrations increase on the membrane surface, a decrease in K_a_^2D^ will cause less complex formation, and we quantify this regime in the Results section. To keep track of these distinct 3D and 2D equilibrium constants, we explicitly retain the 2D superscript for 2D binding, otherwise K_a_ (including K_a_^eff^) describes a 3D constant. To derive a simple analytical expression for K_a_^eff^, we input the pairwise equilibria (Methods) into Eq. 1 and after canceling terms, we use the equilibrium expression

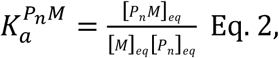

with *n*=1 or 2, to complete the derivation (Methods).

Our main result is then a surprisingly simple and exact analytical relationship that quantifies the equilibrium solution of our model (Fig 1) via K_a_^eff^ and the enhancement relative to K_a_^PP^.

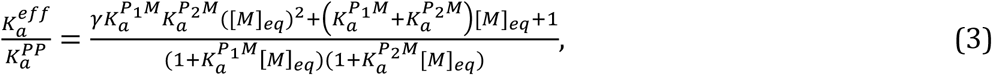

where γ is a dimensionless constant V/(2As), s=K_a_^PP^/2K_a_^2D,PP^, [M]_eq_ is the unbound lipid concentration at equilibrium, and all equilibrium constants (including K_a_^eff^) and concentrations are in volume units. [M]_eq_ is a function of all the model parameters, and can only be solved exactly using numerical methods (Methods); we therefore derive an additional approximate analytical equation for [M]_eq_ described below. However, in the regime where lipids are in excess, the result of Eq. 3 is particularly simple because [M]_eq_∼[M]_0_, the total concentration of lipids. Critically, this means that the initial experimental conditions then directly determine the enhancement. In this regime only two factors control enhancement, the ratio V/(2Aσ), and the dimensionless strengths of membrane localization, 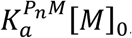 which report the ratio of membrane bound versus solution proteins (Eq. 2) and which we term the membrane stickiness. Hence the volume-to-surface-area ratio, K_a_^PP^/K_a_^2D,PP^, and membrane stickiness play a primary role in triggering (or preventing) constitutive assembly on membranes. In this regime, our Eq. (3) can also be applied to extract binding affinities on membranes (K_a_^2D^) from experiments where binding occurs both on membranes and in solution. This practical application of our result should help simplify the relatively rare experimental characterization of protein-protein affinities on surfaces, as the proteins need not be restricted to the surface for it to work. The condition of excess lipids can be satisfied even with a lipid recruiter such as PI(4,5)P_2_, present at 2.5x10^4^μm^−2^ in the plasma membrane (15), or ∼1% of lipids (13), as we explore further below for proteins involved in CME.

We derive an additional approximate equation for [M]_eq_ to provide a complete equilibrium theory of complex formation applicable to all experimental regimes, and we validate this theory through extensive simulations of ordinary differential equations (ODEs) (Fig 2 and Fig 3, Methods). To briefly outline the derivation, we consider two limiting conditions for localization to the membrane: either there are no protein-protein interactions (K_a_^PP^=0), giving 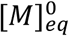, or complete protein-protein complex formation (K_a_^PP^=∞), giving 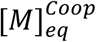 Fig 1d,e). These two bounds are indicated by dashed lines in Fig 2b and both limits are independent of K_a_^PP^. We can continuously interpolate between them (Fig 2b) using the definition

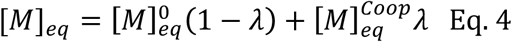

where 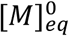 s the root of a quadratic equation, 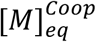 is the root of a cubic equation, and λ is the function of K_a_^PP^ (also the root of a quadratic) that smoothly interpolates between them (see Methods for detailed derivations). This theory then provides a complete description of the equilibrium concentrations of all species, as from K_a_^eff^, one can directly calculate the total complexes formed, and we derive additional equations for breaking these into membrane and solution components (Fig S1, SI Text).

**Figure 2:**
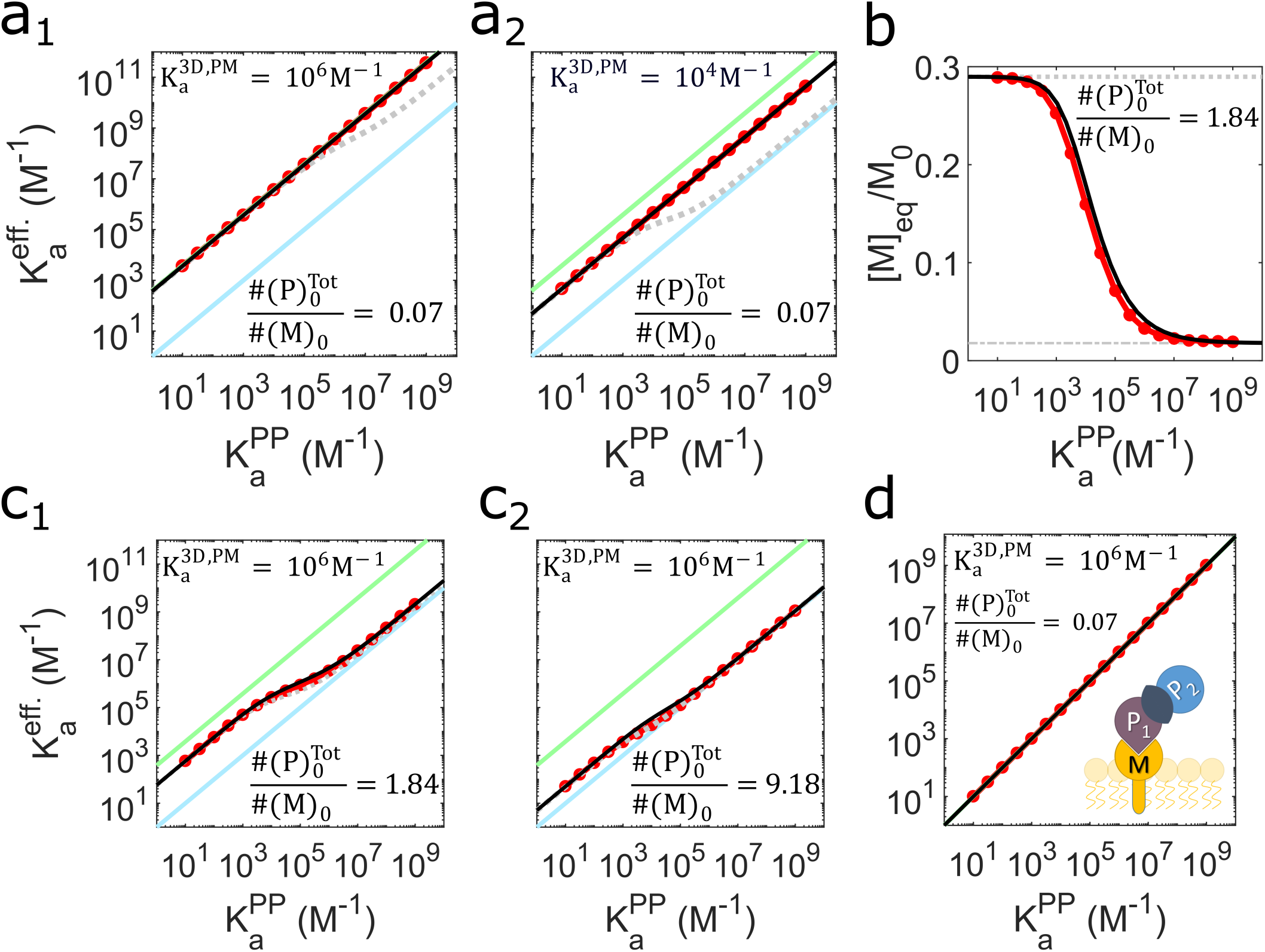
The equilibrium theory developed here accurately predicts how and when membrane localization will enhance protein interaction strength. (a) ODE simulation results shown in red circles are compared with our theory shown in solid black (Eq. 3). For reference, the blue line shows the trend for pure solution binding, i.e. K_a_^eff^=K_a_^PP^. The green line shows the maximum achievable enhancement, occurring if all proteins were on the membrane, K_a_^eff^=γK_a_^PP^. The gray dashed lines are included to contrast the K_a_^eff^ calculated if the cooperative effect is neglected. From a_1_ to a_2_, the K_a_^PM^ is decreased, producing lower but still constant enhancements. (b) The number of unbound lipids is plotted as a function of K_a_^PP^ with all other parameters fixed, showing how lipid binding is a function of the protein interaction strength due to the cooperative effect (Fig 1d,e). The theoretical prediction for [M]_eq_ is shown in solid black. (c) With fewer lipid recruiters relative to total cytosolic proteins ([P]_tot_/[M]_tot_ >1), the enhancement is less pronounced, although for weak binders (low K_a_^PP^) even limited membrane localization causes significant increases in complex formation. From c_1_ to c_2_ the protein concentrations are increased. All results use s=1nm. (d) If only one partner binds the membrane, the protein interaction remains fully 3D and no enhancement occurs.

**Figure 3:**
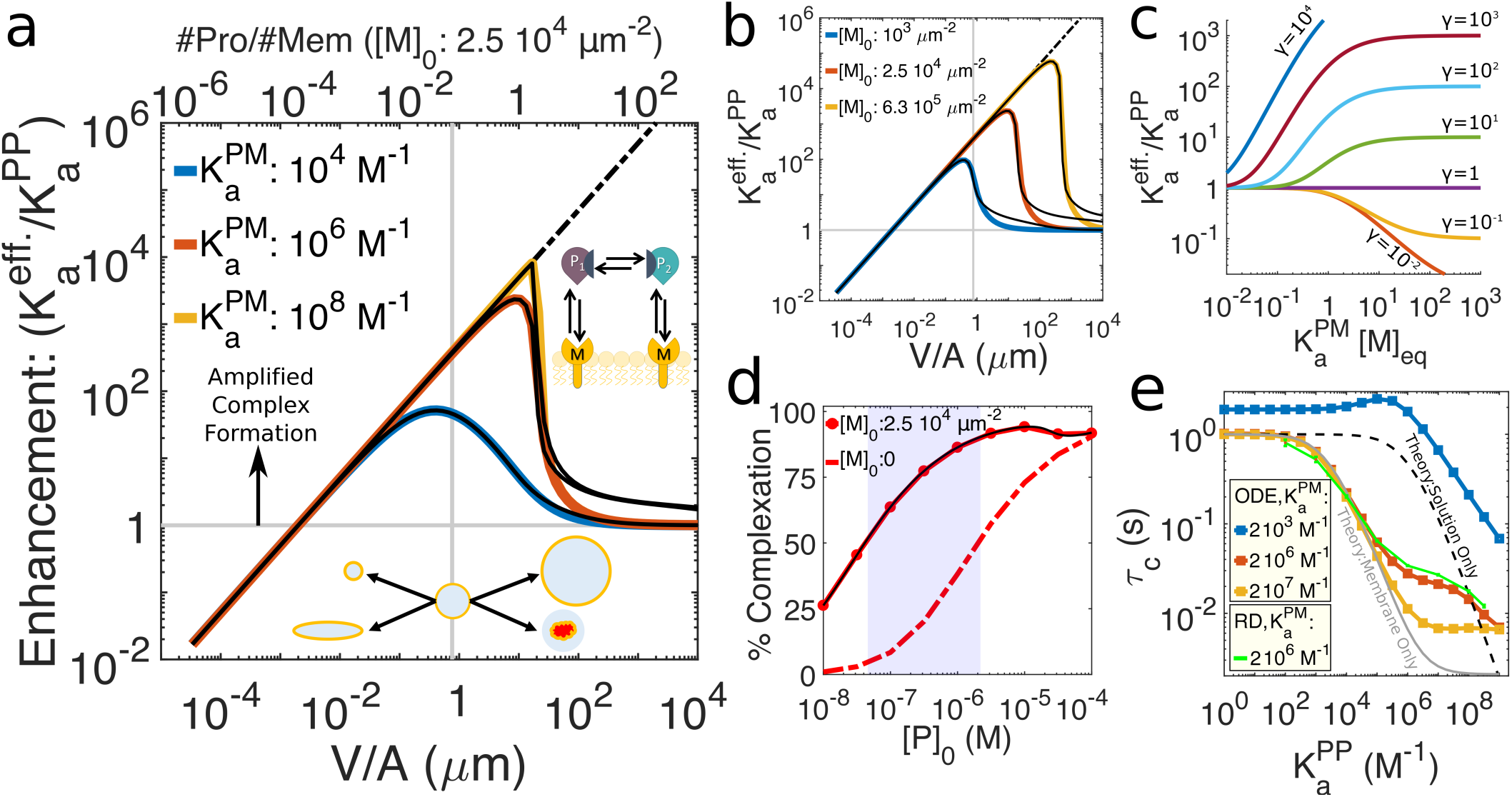
Protein interactions aided by strong protein-lipid interactions, abundant lipid recruiters, and low protein concentrations benefit most widely from membrane localization. a) Enhancement ratios from ODE simulation (colored lines) and theory (black solid lines in all panels), where the upper limit for the enhancement ratio is shown in dashed black line and s= K_a_^PP^/2K_a_^2D,PP^ =1nm. Between the limiting cases where the membrane is either too large to enhance binding (V/A⟶0) or is too small to effect binding (V/A⟶∞), a broad region of enhanced binding occurs. The vertical gray line is the V/A ratio for the yeast plasma membrane, for reference. A maximum for each condition is reached where lipids still outnumber proteins (shown on upper x axis). Increasing (a) protein-lipid affinities and (b) lipid concentrations produces greater possibilities for enhancement. (c) Increases in the membrane stickiness (K_a_^PM^[M]_eq_) produces monotonic increases in enhancement for all values of γ>1. (see Fig S3 for dependence on all other variables). (d) For lower expression levels relative to binding strength (K_a_^PP^=10^6^M^−1^),membrane localization can act as a switch to turn on assembly from <50% to >50% (shaded areas). (e) Timescales to equilibrate were calculated from simulations of both ODEs (solid lines) and reaction-diffusion (RD) (green points) (Methods). Weak lipid binding can reduce speeds (blue) relative to pure solution binding (dashed). The approximate theoretical bounds shown here for time-scales of binding either purely in solution (dashed) or on the membrane (gray) derive from the kinetics of irreversible association (SI Text).

## RESULTS AND DISCUSSION

### Bounds on binding enhancement are determined by V/A and Ka^PP^/Ka^2D,PP^

Using our main result, Eq.3, one can predict when and by how much membrane binding will enhance complex formation of binding pairs without performing any simulation or experiment. To establish possible values for K_a_^eff^, we first ask: are there any cases where membrane binding will reduce protein complexation, i.e. K_a_^eff^<K_a_^PP^? To answer this, we consider the case where all proteins are on the membrane 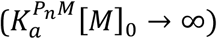, such that we have pure 2D binding and K_a_^eff^=γK_a_^PP^=K_a_^2D,PP^ V/A. Reduced protein complexation will occur only if γ<1, or V/A<2σ. The size of s controls the relative strength of K_a_^2D,PP^ vs. K_a_^PP^ and varies from one binding pair to another, but experiment and theory indicate it is of the nanometer length-scale (21, 22, 25). We therefore collected the V/A ratios for a wide range of cell types and organelles to illustrate that in nearly all cells, V/A>2σ and thus membrane localization will enhance binding (Fig S2). Indeed, targeting the plasma membrane in most cells results in γ values much greater than 1, in the range 10-1000 (Fig S2). Ultimately, the V/(Aσ) ratio is absolutely central in controlling observed enhancement, as it sets the maximum achievable K_a_^eff^. In most cases proteins will end up mixed between solution and membrane, and from Eq. 3 this gives us K_a_^PP^≤K_a_^eff^<γK_a_^PP^ (Fig 2). In the cases where only one protein binds to lipids, all 2D localization benefits are lost and no enhancement occurs: K_a_^eff^=K_a_^PP^ (Fig 2e).

### Cooperativity emerges in protein-lipid binding

An important feature that our analysis captures is the coupling that emerges between protein-protein affinity and protein-lipid binding due to membrane localization. If a protein is localized to the membrane by a partner, binding to a second lipid then becomes a 2D rather than a 3D search (12) (Fig 1d,e). Thus, stabilization of proteins on the membrane is achieved not only through strong protein-lipid interactions, but by feedback from strong protein-protein interactions. This cooperative effect for lipid binding produces the unexpected result that the number of proteins bound to the membrane is dependent on the protein-protein interaction strength (Fig 2b). If this cooperative effect is ignored, K_a_^eff^ is underestimated, particularly for strong binding proteins (Fig 2, gray dashed lines, SI Text). Our theory (Eq. 3) and our equation for [M]_eq_ (Fig 2b) explicitly capture this important cooperative effect.

### Anticipating binding enhancement and response to factors external to the model

With our theory, it is possible to directly probe how changes in cell geometry, binding affinities, or concentrations will regulate enhancement. Although the model is too simple to fully describe multi-component assembly, by varying the input parameters to mimic competing cytosolic factors, one can evaluate the relative importance of concentration, affinities, and geometry. This is particularly true of equilibrium *in vitro* experiments. For changes to geometry, we first note that the equilibrium results depend only on the ratio and not the absolute size of V or A, and the membrane does not have to surround the solution volume like the plasma membrane but can reflect binding to the outsides of liposomes, for example. Although it may seem that increasing V/A (through γ in Eq. 3) always increases K_a_^eff^, this is not the case when [M]_0_ is kept constant in its natural units of μm-2 (it is then converted into Volume units by multiplication by A/V). V and A thus also control the initial copy numbers of proteins and lipids separately, such that large values of V/A have a great excess of proteins over lipids. This drives K_a_^eff^⟶K_a_^PP^ (Fig 3a,b). For most physiologic V/A values (∼0.05-20μm Table S3) however, and physiologic concentrations of proteins (1nM-10μM) or lipids (103-105μm-2 Table S4), proteins are not in great excess, meaning significant enhancement is achievable depending on the membrane localization strength (Fig 3a,b).

#### Membrane composition

Because the membrane is described in our model only by its surface area A and the concentration of lipids targeted by the protein lipid-binding domains, the spatial and chemical heterogeneity of cell membranes is not captured. The stickiness of a membrane for specific proteins is captured in our model, however, by the dimensionless product K_a_^PM^[M]_eq_, which is simply equivalent to one protein’s ratio of membrane bound to unbound copies (e.g. [MP_1_]/[P_1_]) at equilibrium. As is clear from Fig 3c, the enhancement is a monotonically increasing function of K_a_^PM^[M]_eq_ for all γ>1 geometries. Hence with either stronger affinity for the membrane (K_a_^PM^) or a higher concentration of target lipids ([M]_0_), localization to the membrane and thus enhancement will increase. Fig 3c also illustrates how once K_a_^PM^[M]_eq_ exceeds values of ∼10, maximum enhancement is reached and increasing the stickiness of the membrane does not change the resulting binding. In the SI text, we derive an explicit formula for this critical value of K_a_^PM^[M]_eq_, as well as the corresponding critical value for lipid concentration, [M]_c_ (Fig S3a,b). If the lipid concentration and affinity contribution to stickiness can be deconvoluted, as is the case for proteins that target individual lipids (such as PI(4,5)P_2_) at essentially a 1:1 ratio, one can individually evaluate how decreases in either lipid concentration (Fig 3b) or affinity (Fig 3a) will lower enhancement. Importantly, we find that surprisingly low lipid concentrations are sufficient to drive maximal enhancement. For V/A=1μm, (about the value for yeast cells), the PI(4,5)P_2_ concentration of 2.5x104μm-2 produces close to the maximum in enhancement, meaning adding more lipids makes minimal difference (Fig 3b, Fig S3ab). For proteins such as BAR domains, in contrast, assigning values for affinity and lipid concentration would require a composition dependent interpretation of these values, as BAR domains are less selective for individual lipid types and may only bind stably to clusters of lipids rather than 1:1 (see SI Text for extended discussion). Nonetheless, any change in membrane composition that increases its stickiness towards any specific membrane binding domains will clearly drive up binding interactions between associated protein pairs (Fig 3c).

#### Competition for protein and lipid binding

Thus far we have said little about protein concentrations or K_a_^PP^, as enhancement is independent of their magnitude when lipids are in excess. However, these protein variables always determine when enhancement acts as a switch to turn on complex formation. Some proteins are perfectly capable of forming strong complexes in solution, whereas protein pairs with less than 50% complexes formed in solution can experience dramatic increases in bound complexes (Fig 3d, Fig S3e). The dependence of enhancement on protein concentration is also monotonic for fixed geometries, whereas protein concentration drops, enhancement increases (Fig S3d). Hence, competition for binding any of our protein binding pairs in solution would be expected to lower initial concentrations of each component. This will increase the ultimate enhancement (Fig S3d) and in many cases, make the proteins more sensitive to localization as a trigger for assembly (Fig 3d). If competition for protein binding partners involved other proteins that also bound to the membrane, then enhancement could be increased, but this extension beyond the Figure 1 model would have to be quantified via simulation. Competition for lipid pools, on the other hand, will always decrease enhancement, as shown in Fig 3b and explored further below for CME proteins.

#### Sensitivity to protein-protein affinity

Mutations to proteins would largely affect their affinities, either for their protein or their lipid partners. As noted above, however, even many-fold changes in affinity may have minimal consequences on measured enhancement (Fig S3g). For mutations to ENTH/ANTH domain containing proteins, a sufficient concentration of target PI(4,5)P_2_ lipids prevents any significant change in enhancement despite 10-fold changes in K_a_^PM^(26) (Fig S3h). Decreases in K_a_^PP^ can have similar effects, either not affecting enhancement when membrane stickiness is already high, or otherwise increasing enhancement (Fig S3c, Fig 2). For complex formation, decreases in K_a_^PP^ can be much more significant, acting to increase the sensitivity of complex formation to localization, such that it is more likely to act as a trigger for assembly (Fig S3e). Mutations that asymmetrically affect the K_a_^2D^ values would result in changes in σ values, with smaller values always driving larger enhancement and stronger complexation on the membrane (Fig S3f).

### Timescales of assembly vary strongly with binding affinity

Our theory only describes the equilibrium state of the model, but we can determine speeds of assembly via simulation. Now the binding rates and the absolute values of V and A (not just the ratio) will influence the kinetics (all simulation inputs in Datasets 2, 3 and 4). For these time-scales, we find that protein-lipid affinities K_a_^PM^ are most often shown to be critical in controlling the overall time-scales of complexation, even driving slow-downs in speeds relative to solution binding (Fig 3e, Fig S4). Changes in diffusion from solution to the membrane (about 100 times slower) affect the magnitude of association and dissociation rates and are captured implicitly in our ODE simulations (Methods), and explicitly in our spatially resolved reaction-diffusion simulations (19, 20). However, the influence of diffusion on the reaction rates is rarely a dominant factor in physiological rate regimes (Fig S4), indicating it is the binding strengths rather than slow 2D diffusion that determine assembly speeds. However, we note that our comparison of ODE and RD kinetics was performed in relatively small systems, and it is true that as spatial dimensions increase, times to diffuse to reach the membrane will influence the overall equilibration times. The timescales we calculated for protein pairs and scaffold mediated systems (Figure S4b, c) were performed using ODEs at their corresponding cellular dimensions (Table S3): V=1200 μm3 (human) and V=37.2 μm3 (yeast). Using RD simulations would produce slower relaxation times, particularly for human cells, due to the time required to reach the surface. Crowding would also lower effective diffusion constants of proteins, although the decrease in time-scales to equilibrate would be negligible unless binding rates were strongly diffusion-influenced (Methods).

### Biological relevance for proteins in CME

To test the biological relevance of membrane localization for driving complex formation and assembly, we collected biochemical (Table 1, Table S1-S2), concentration (Table 1, Table S4), and cellular geometry data (Table S3) for interactions among 38 membrane targeting proteins in yeast and human cells, including 22 proteins involved in clathrin--mediated endocytosis (CME). We first study only individual protein pairs that can bind according to our model of Figure 1, (Table S1) shown in Fig 4a: the membrane binding proteins AP-2, DAB2, ARH, FCHo1, FCHo2, HIP1, HIP1R, PICALM, SH3GL2, EPN, AP180, SLA2, and SYP1. In Figure 4b,c we show results of binding between specific pairs. We used cytosplasmic concentrations of the proteins (Table 1) and the targeted lipids (Table S4), and the relative Volume and Area from their respective cell types (Table S3). Binding constants are collected from previous experimental studies (Table 1, Table S1, Dataset S3), and for 2D binding constants we test values of s=K_a_^PP^/2K_a_^2D,PP^=1nm (Fig 4) and 10nm (Fig S5). Our results thus provide quantitative insight into how these pairs in isolation would use membrane localization at physiologic conditions to drive their protein-protein interactions. For some proteins, such as AP-2, solution 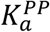 values have been measured with partners (Fig 4a, Table S1), but further experiments indicate that the proteins undergo minimal binding in solution due to conformational regulation (1). Despite this additional regulation, membrane localization will still increase complex formation relative to what is observed in solution (γ>1), so the effect is quantified here using the measured 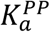 value. Using our theory along with simulations for verification and time-scales, we find that affinities of these CME binding pairs can be enhanced 10-1000 fold by binding to membranes (Fig 4b, Fig S5 for results with σ=10nm). With limited binding in solution for most pairs, membrane localization then triggers a dramatic increase in complex formation (Fig 4c, Fig S5). The central adaptor protein AP-2 is responsible for many of these interactions, showing the capacity to trigger assembly with nearly all of its binding partners (Fig 4c, green bars). Even though we assume binding is possible for AP-2 in solution, it is still quite limited prior to localization. Not surprisingly, knockdown of AP-2 in mammalian cells causes severe disruption of endocytosis (27), underlining its secondary importance only to the irreplaceable clathrin (27) and PI(4,5)P_2_ (28). We note that AP-2 can potentially bind up to three PI(4,5)P_2_ copies, meaning that there will be less free lipids available for each AP-2. With fewer lipids, enhancement and complexation will be reduced, but is still quite large (Fig S5). For some proteins such as FCHo1 (SYP1 in yeast), binding affinities (K_a_^PP^ or K_a_^PM^) are not available, and this F-BAR protein does not target a single lipid specifically. However, by considering ranges of membrane stickiness values, we can use our method to identify which combinations (Fig S6) best describe the experimental observation that these proteins only localize effectively to membranes when they can bind other proteins (18, 29). We find for this protein, membrane stickiness values of ∼0.5 produce membrane targeting that is sensitive to protein-protein interactions, whereas once values exceed ∼1, no partners are needed to target the membrane effectively (Fig S6).

**Figure 4:**
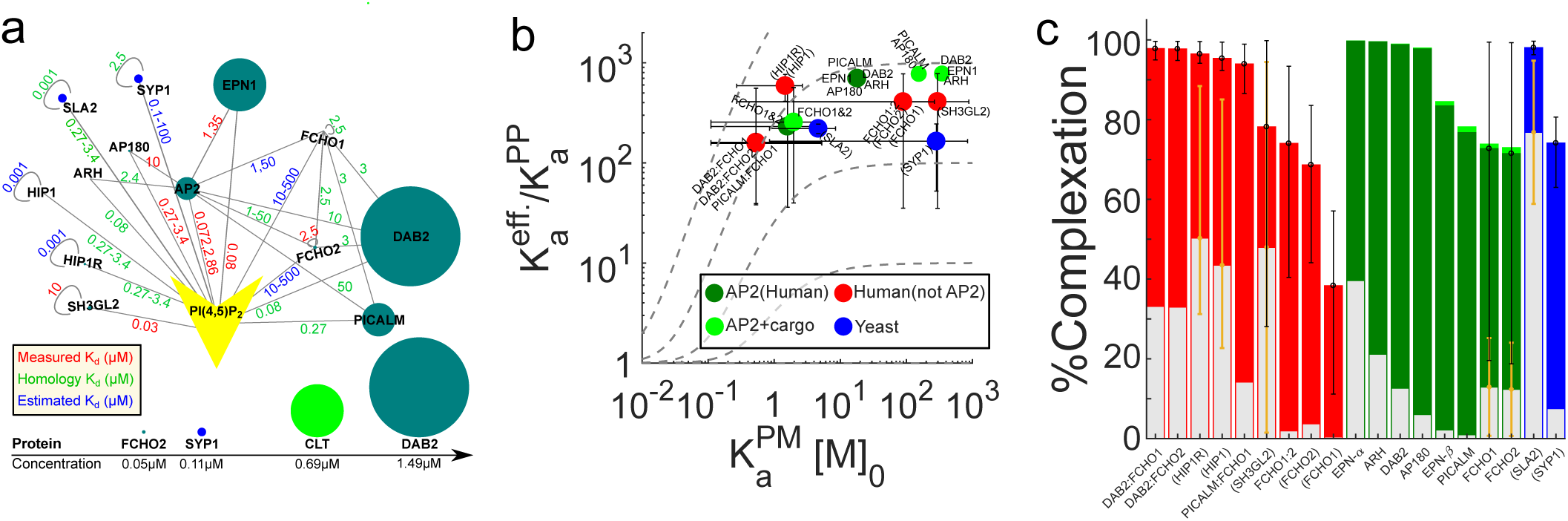
Membrane localization triggers strong complex formation for pairs of protein binding partners involved in clathrin-mediated endocytosis. (a) Interactions between human (green) and yeast (blue) CME proteins that also bind PI(4,5)P_2_ are shown along with K_d_^PP^ values (μM) either measured (red text), inferred through structural and functional homology (green), or estimated (blue) (Table 1, Table S1, Dataset S3). Sizes indicate concentrations (Table 1). (b) Enhancements for each of these CME binding pairs, following the model of Figure 1, were calculated using Eq. 3 and verified through numerical simulation of ODEs (Dataset S3). Pairs involving human AP-2 are in dark green, light green points involve AP-2 with cargo binding-adjusted K_a_^PM^, with red and blue points showing other human and yeast proteins, respectively. The x-axis is the average membrane stickiness for each pair, and here we assume s=1nm (see Fig S5 for s=10nm). For poorly characterized lipid binding affinities, we considered ranges of values (error bars in x), producing ranges of enhancements (error bars in y). Gray lines are guides for fixed V/A ratios. Protein names in parentheses are homo-dimers. (c) The percent of proteins in complexes increases from solution binding levels (gray bars) as a result of membrane localization (colored bars). All results and parameters used for all data points in Dataset S3.

We further interrogate two additional mechanisms for stabilization at the membrane by lipid binding proteins such as AP-2, epsin, and Dab2 (3). First, they each bind transmembrane cargo after membrane localization, which acts to effectively increase the K_a_^PM^ by increasing their residence time on the membrane. K_a_^PM^ is a factor of ∼40 higher for AP-2 binding to PI(4,5)P_2_ when cargo is available (30). Interestingly, these cargo stabilized interactions (Fig 4b, light green) do not make a significant impact on complexation when we assume the full 1% PI(4,5)P_2_ concentration is free to bind, as the numerous lipids outweigh a need for stronger binding (Fig 4c, light green). However, when we evaluate complexation with PI(4,5)P_2_ pools diminished by a factor of 10 due to assumed competition from other PI(4,5)P_2_ binders, now cargo stabilization via higher K_a_^PM^ does help recover strong complexation on the membrane (Fig S5). This suggests that cargo binding becomes functionally relevant for stabilization when competition from multiple adaptors limits PI(4,5)P_2_ binding. Second, when these adaptors can bind multiple partners with distinct appendage domains (31), we see more proteins on the membrane due to the increased difficulty of un-tethering from the membrane domain (Fig S6b).

In the cellular environment, CME proteins are of course not in isolation and can both compete and cooperate with one another to form higher order assemblies, induce conformational changes, and occupy lipid binding sites on the membrane. Thus we can only speculate about the role of localization in nucleating clathrin-coated pits *in vivo.* It is undeniably true, however, that by increasing the concentrations of binding partners without a dramatic drop in K_a_^PP,2D^ relative to K_a_^PP^, which is what occurs in physiologic cases as shown above, proteins will increase complex formation on the membrane. The initial nucleation of clathrin-coated pit sites is difficult to resolve experimentally because of the challenges in tracking the many participatory proteins simultaneously, and because prior to cage formation, the density of molecules is, by definition,low. Experiments have tracked the role of AP-2 and clathrin in nucleating sites (32), which we discuss below.

### Scaffold mediated interactions of CME proteins also exploit membrane localization

To go beyond our Figure 1 model of pairwise protein binding and thus characterize how scaffold proteins (Table 1: ITSN1, EPS15, EDE1 and SLA2) stabilize complex formation at the membrane despite not directly interacting with the lipids (model in Fig S7, list of interactions in Table S2), we simulate systems of ODEs (Methods). We thus simulate interactions involving three proteins, two of which can bind lipids but not each other, and the third that binds both peripheral membrane proteins but not the membrane (Fig 5a). Our results in Figure 5b,c show that while scaffold mediated complexes can still capitalize on 2D localization for binding (Fig 5b), because localization is now mediated by peripheral membrane proteins that are at much lower concentrations than the lipid recruiters, we find that the increase in complex formation is less robust (Fig 5c). Results are now also sensitive to the concentration of the scaffold protein (Fig S8a), as without scaffold proteins, no localization benefits or enhancement is possible.

**Figure 5:**
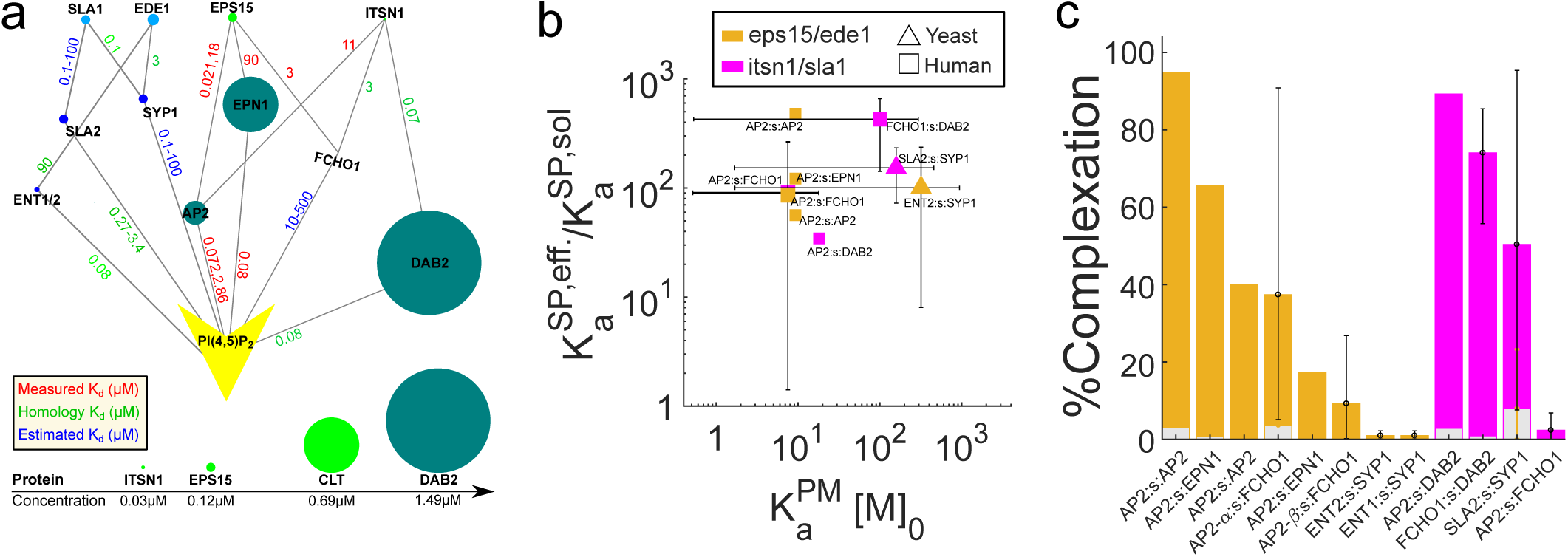
Scaffold-mediated interactions can also exploit membrane localization to drive complex formation in clathrin-mediated endocytosis. a) When sets of three proteins can form a complex and two of them also bind to lipids, localization is again capable of driving stronger complex formation. Yeast proteins in blue and human proteins in green, where the uppermost four (SLA1, EDE1, EPS15, ITSN1) are scaffold proteins that do not bind lipids. Affinities follow the same color scheme as Figure 4, and we have included only binding interactions between these proteins that are shown in parts b-c. (b) Because our primary model (Figure 1) no longer applies, we cannot use Eq. 3 to predict enhancements, and instead use simulations of ODEs. The scaffold model is defined in SI Text and Figure S7, along with the definitions of K_a_^eff,SP^ and K_a_^sol,SP^, where neither is a true equilibrium constant and the equilibrium results of the simulations are used to calculate their defined values. Without membrane localization, K_a_^eff,SP^→K_a_^sol,SP^. Scaffold proteins (*s* in the labels) are eps15/ede1 (orange), or itsn1/sla1 (pink). (c) Complexation involves all three proteins (see SI Text) and here again, the percent of proteins in complexes increases from solution binding levels (gray bars) as a result of membrane localization (colored bars). All results and parameters in Dataset S4.

### Clathrin cage nucleation and BAR domain oligomerization can exploit membrane localization

Thus far we have not discussed clathrin, the central component of the CME vesicles that does not actually bind to lipids itself. *In vitro* experiments find that clathrin polymerization on the membrane (via adaptor binding) is more robust than occurs in solution (with adaptors still present), supporting a role for membrane localization in its nucleation and assembly (2). Clathrin is a trimeric protein with three binding sites to target peripheral membrane proteins. It polymerizes with itself into hexagonal lattices without competition from the peripheral membrane proteins. Thus, its interactions with peripheral membrane proteins not only increase the quantity of protein bound to the membrane, it can help drive 2D polymerization between clathrin trimers. Through (non-spatial) stochastic simulations (Methods) set-up to mimic recent *in vitro* experiments (1), we explored a range of clathrin-clathrin interaction strengths to show how membrane recruitment by the AP-2 adaptor (1) can enhance clathrin polymerization yield (Fig S8). We find that clathrin localizes to the membrane first via AP-2 binding before assembling into cages in 2D for the most reasonable K_d_s of 10-100μM (33). This result is supported by evidence from *in vivo* experiments that probe the early stages in the nucleation of clathrin coated pit sites through tracking of AP-2 and clathrin (32). They found that clathrin arrives at the membrane most frequently (75%) as a single trimer, and bound to at least one but most often two AP-2 molecules (32). Nucleation can then proceed in two ways: 1) another clathrin trimer localizes to the membrane via AP-2 and these trimers dimerize in 2D or 2) another clathrin trimer is directly recruited by the clathrin on the surface. Although the subsequent clathrin dimerization events were not resolved experimentally, preventing definitive evidence of membrane localized clathrin-clathrin assembly, the fact that each clathrin is bound to AP-2 suggests that solution clathrin recruits to the membrane via the first pathway: AP-2 binding to lipids. From our simulations, this first pathway is markedly dominant. There is a strong driving force both from affinity and from concentration for AP-2 to bind any of the 25000 PI(4,5)P_2_/um^2^, and correspondingly minimal drive for a solution clathrin to bind a small number of clathrin trimers localized to the surface. We note that because clathrin also arrives at the membrane as dimers or higher order complexes 25% of the time, solution binding of clathrin also contributes to nucleation of pit sites, but to a much lower extent (32). Interestingly, once pit sites have formed, assembled clathrin cages exchange with solution clathrin with the aid of ATP-consuming proteins that facilitate remodeling of the clathrin cage (34). Thus clathrin-clathrin interactions from solution certainly play an important role in the cell in maturing the pit sites (34).

CME proteins with BAR domains that dimerize, appear to oligomerize only on the membrane, and are functionally important for driving membrane deformation (17, 18) can also exploit localization to drive their binding interactions. We study isolated FCHO1/2 oligomerization and endophilin (SH3GL2) oligomerization (Fig S9). Here again we consider a range of K_a_^PM^ values to capture uncertainty in the membrane stickiness of these domains. We find the stoichiometry of the dimerization pair (homo or hetero) is central in determining whether large oligomers form. With matched pairs, homodimers such as endophilin form larger oligomers that feedback into higher stabilization at the membrane, whereas the disparity in FCHo1 and FCHo2 concentration (Table S1) produces more isolated dimers. Experiments have shown that BAR domains exhibit stronger binding to curved membranes (17). Because we lack this cooperative feedback in our model between oligomers tubulating membranes and thus potentially increasing affinity for subsequent proteins, our result can be interpreted as a lower bound on observed oligomerization.

In all cases, an important outcome of these strong binding interactions on the membrane is that they are difficult to disassemble, consistent with findings that unproductive assembly events observed *in vivo* (35) require the ATP-driven uncoating machinery for disassembly (2, 36). Our results demonstrate that establishing the physiologic significance of these polymerization observations depends not only on protein concentrations and solution conditions, but also the V/A ratio. Thus, this ratio should be regarded as a critical factor in comparing *in vitro* and *in vivo* behavior.

### Biological relevance for membrane remodeling pathways in yeast

Lastly, our analysis reveals that functionally diverse proteins that target membranes in yeast can follow pathways to assembly both similar and distinct from the CME proteins. In particular, the CME protein pairs produce limited protein-protein complexes when isolated in solution, but experience large enhancements due to membrane localization, triggering widespread protein-protein interactions only after binding to the membrane (Fig 4). For 15 yeast proteins involved in daughter cell budding, lipid regulation, and morphogenesis, we studied their pairwise binding in using cytoplasmic (yeast) concentrations (Table 1), lipid concentrations (Table S4), cytoplasmic V/A ratios (Table S3) and experimentally measured protein-lipid affinities (Table 1). We find binding enhancements are high (100-1000), similar to the CME proteins, indicating that binding will be promoted once proteins are on the membrane (Fig S10, Dataset S2). Although enhancements were readily measured for these yeast binding pairs because they were independent of K_a_^PP^ values (Dataset S2), we could not directly compare complexation for these interactions as we did for the CME interactions because they lacked any K_a_^PP^ data. For binding enhancements, we found an exception in the coat forming proteins targeting endosomes (VPS5, VPS17, SNX4, SNX41), which only exhibit enhancements <20. These proteins target the PI(3)P lipid but most bind only weakly (K_d_^PM^>100μM) (37), limiting their enhancements despite a favorable V/A ratio at the endosome (Fig S10, Table S3). Unlike in CME, however, these coat proteins form stable interactions in solution (38). Thus, rather than membrane binding triggering protein interactions, we would first expect the reverse: strong protein interactions in solution function to target and stabilize protein at the membrane through the cooperative effect (Fig 1d,e, Fig 2b). We test how the binding of the retromer components VPS5 and VPS17 to the endosome will be significantly enhanced by forming a higher order assembly in solution with the strong lipid binding cargo adaptor, SNX3 (Fig S9). SNX3 targets PI(3)P with stronger affinity (∼2μM) than either VPS5 or VPS17 (37), and is known to improve recruitment of the retromer to endosomes (38). Once these small pre-assembled coat subunits are on the membrane, they can then continue to exploit localization to form larger protein coats.

## CONCLUSIONS

We conclude by noting that assembly on membranes is regulated to occur at specific times or sub-cellular locales, and our theory provides a useful aid in predicting the changes in local protein, lipid concentrations, and affinities that are necessary to trigger (or prevent) such assembly. Ultimately, our theory is most powerfully applied to interpreting *in vitro* results, due to the simplifying assumptions of the model, and can improve the design and quantitative interpretation of assays probing multi-protein complexation at membrane surfaces. Also, given known protein-protein and protein-lipid binding affinities, our theory can quantitatively predict the results of *in vitro* experiments that mimic Figure 1, thus avoiding the need for such measurements. Our results indicate that even relatively low lipid concentrations (i.e. PI(4,5)P_2_ at ∼1% of plasma membrane lipids) can be sufficient in many cases to stabilize proteins to membranes and drive protein-protein interactions. We found that additional factors, such as cargo binding by adaptor proteins in CME, are only strong regulators of membrane localization or protein interactions under specific conditions. Cargo-binding became significantly more important when we reduced total PI(4,5)P_2_ concentration to mimic the physiologically relevant condition where much of the lipid may be unavailable due to competition from other proteins. A fruitful means of exploring in more detail the role of cytoplasmic factors, as well as spatial heterogeneity, crowding, and non-equilibrium dynamics, is through reaction-diffusion simulations, although we note the results will then be dependent on many additional parameters. Overall, the theory we provide here offers a general and useful quantitative guide for predicting when or if membrane localization plays a role in the cellular control of self-assembly.

## METHODS

### Theoretical Derivations

**A1. *Derivation details of K*_*a*_^*eff*^ *(Eq. 3)*** The exact solution (both equilibrium and time-dependence) to our model of proteins interacting and recruiting to membranes (Fig 1) can only be obtained numerically. Starting from our definition in Eq. (1), 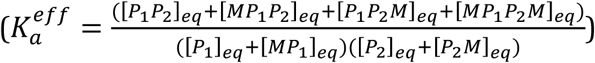

We input pairwise equilibrium expressions for each species In the numerator, where the complete list of pairwise equilibria illustrated in Fig 1 are given by equations:

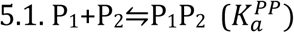

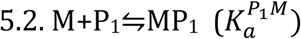

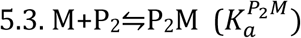

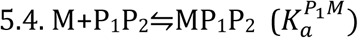

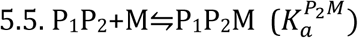

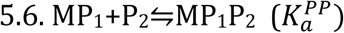

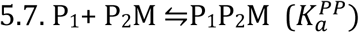

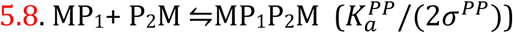

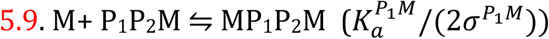

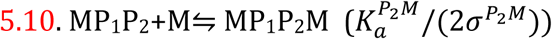

where 5.8, 5.9 and 5.10 are all in 2D. Reactions in 2D list the 2D K_a_ values and thus require species be in units of Area-1. To solve all species in consistent units, where we will use solution concentrations with units V^−1^, the listed 2D K_a_ values must be multiplied by V/A.Inputting Eqs. 5.1, 5.6, 5.7, and 5.8 into the numerator of Eq. 1 and dividing numerator and denominator by the factor [P_1_]_eq_[P_2_]_eq_ gives:

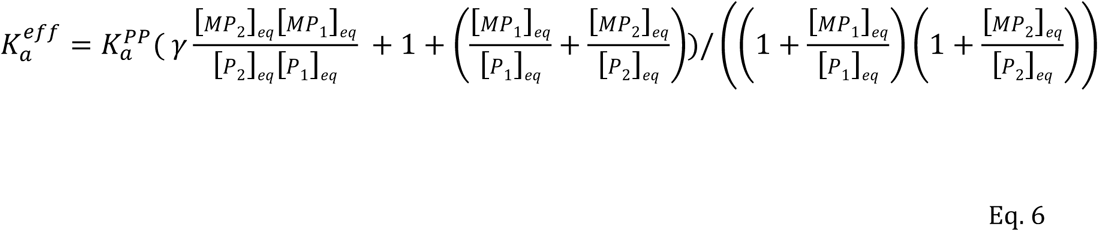

where 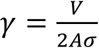 And finally using Eqs. 5.2 and 5.3 above (Eq. 2 of the main text),we recover our main result, Eq. 3. We also note if P_1_ and P_2_ target distinct lipids, the [M] concentrations will be subscripted accordingly. Thus Eq. 3 of the main text is exact. However, a separate equation for [M]_eq_ is needed that will be approximate.

**A2. *Derivation details of [M]*_*eq*_ *(Eq. 4)*** Our equation for the unbound lipids at equilibrium, [M]_eq_, is a function of two limiting cases for protein localization to the membrane with a smooth interpolation in between defined via Eq. 4 of the main text 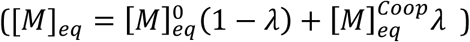 in the first extreme (K_a_^PP^=0), we solve for unbound lipids at equilibrium based solely on protein-lipid interactions, M+P⇌MP, giving the familiar quadratic root

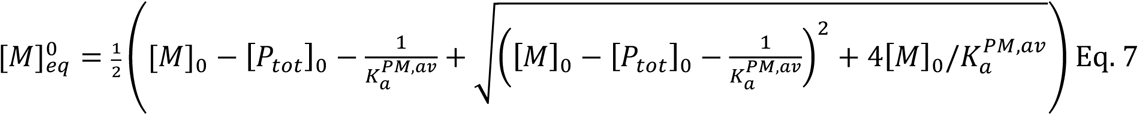

This equation recovers [M]_eq_∼[M]_0_, the initial concentration of lipids, when [M]_0_>>[P_tot_]_0_=[P_1_]_0_+[P_2_]_0_. The K_a_^PM,av^ is the average from both proteins. For significant differences between affinities and protein populations, the weighted average is most accurate, 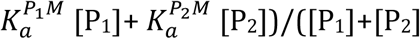, where using [P_1_]_eq_’s rather than [P_1_]_0_ is more accurate. For the second extreme, (K_a_^PP^=∞) we now treat all proteins as bound in complex, creating an equilibrium between proteins with two lipid binding sites and the membrane. This gives us 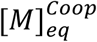, which has a cubic root analytical solution (see SI Text). Lastly, we interpolate between these two extremes (both independent of K_a_^PP^), using the function λ. The λ function is the fraction of proteins that are bound to one another (based on P_1_+P_2_⇌P_1_P_2_ with K_a_^eff^) out of the maximum possible,

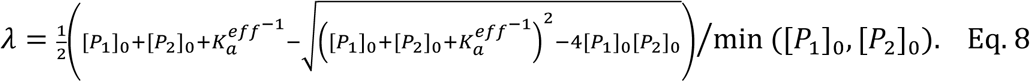

These three results thus complete the equilibrium theory for Eqs. 3-4.

An important feature of Eq. 4 is that it produces the correct limiting behavior as λ goes from 0 to 1. We emphasize that although the equation for 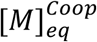 s cumbersome, simply setting λ=0 will already give very good accuracy in reproducing the exact result, with noticeable errors only expected when both the lipid concentration is small relative to the total proteins and the K_a_^PP^ is large. We note that the relative error in the full Eq. 4 is quite small, although for large V/A ratios (larger than those observed physiologically), the error grows and produces overestimates of theoretical enhancement ratios relative to the numerical solution (Fig 3a,b). We provide an excel workbook that performs this complete calculation for any model, including if each protein targets distinct lipids.

### B. Definition of microscopic and macroscopic rates for simulation

To simulate the systems of ordinary differential equations (ODEs) for Fig 1 (Fig S1e, SI Text), we need macroscopic rates, and to simulate the single-particle reaction-diffusion system (RD), we need microscopic rates (also known as intrinsic rates in the Smoluchowski theory (39)) in both 3D and 2D. The macroscopic rates emerge based on the dynamics of the more detailed microscopic system, and can therefore be constructed to optimally match the kinetics of the ODE simulations to the RD simulations. We note that these definitions are specific to the kinetics, as the equilibrium of both simulation approaches will be identical due to their matching equilibrium constants.

The ODE simulations do not account for space or explicit diffusion. Here, we define their rates to implicitly account for changes to diffusion and thus best match the RD simulation kinetics. Macroscopic association (on-) rates can be defined in 3D from the intrinsic rate of the Smoluchowski model via the relation (40):

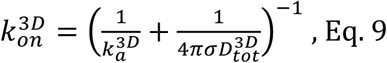

where k_a_ is the intrinsic association rate that captures the barrier to complex formation for species in contact at binding radius s, and D_tot_ is the sum of both species diffusion constants. The macroscopic off-rate can be defined in all dimensions via

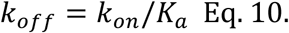

The intrinsic dissociation rate k_b_ is defined via the corresponding equation, k_b_=k_a_/K_a_, with all off-rates having the same units in all dimensions of s-_1_. In 2D, there is no single macroscopic rate constant independent of the system size or concentrations (20). However, one can define a macroscopic 2D rate, built on theory from Szabo et al (41), that provides optimal agreement with the corresponding spatial reaction-diffusion simulations via (20):

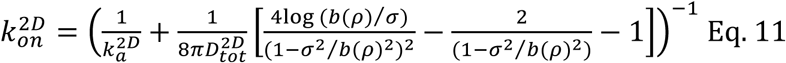

Where

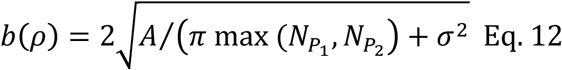

is a length scale that is defined based on the more concentrated of the reacting species P_1_ or P_2_ in the surface area *A*.

The important interpretation of Eqs. 9 and 11 is that, unless k_a_ is large, even substantial (factor of 10 or more) changes to the diffusion constant will have a relatively small impact on the macroscopic rate. It is not until macroscopic rates reach values of ∼106-107M-1s-1 that they become strongly diffusion influenced and thus sensitive to changes in diffusion.

Our 2D intrinsic rates are defined relative to our 3D rates via

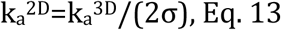

and unbinding rates

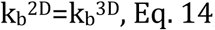

which produces the equilibrium relation defined in the main text, K_a_^2D^=K_a_^3D^/(2σ). We assume here that the dissociation rates are the same from 3D to 2D. It is the association rates that capture two species finding one another in a specific spatial dimension. This definition of Eq. 13 also can be shown to preserve the reactivity of the binding interaction in the Smoluchowski model from 3D to 2D, independent of changes to diffusion (SI Text). For the macroscopic 2D rates, k_on_^2D^, we used Eq. 13 in Eq. 11, which allows us to capture effects of diffusion towards timescales of binding in k_on_^2D^, as D2D is ∼100 times lower than D3D. Transitioning from solution to the membrane via binding lipid or protein involves a 3D search, and thus uses the corresponding 3D rates. See SI Text for further discussion.

Ultimately, the results of K_a_^eff^ are only sensitive to equilibrium constants such as K_a_^2D^ and therefore the size of σ, rather than sizes of relative rates. This length scale σ encodes thermodynamic properties of the molecules involved in the binding reaction and is of the nanometer range (22). In general, the value of σ therefore depends on the proteins involved, but s (or K_a_^2D^), is almost never measured. We extract σ∼7nm (from V/A=6.7μm and K_a_^eff^/K_a_^PP^∼500) in the experimental measurement of 2D binding between calmodulin and a target peptide (21).

Smaller σ values have been observed (25). For simulations, we thus used either 1 or 10nm, showing the expected dependence in Fig. S3f. We used the same value for the protein-protein (s_PP_) or protein-lipid (s_PM_) 2D binding interactions, although only σ_PP_ appears in Eq. 3. The size of σ_PM_ is constrained to ensure an equilibrium steady-state is reached, such that s_PM_ = s_PP_.

### C. Computer simulation methods

**C1. *Numerical solutions of ODEs*** The majority of our simulation results (exceptions noted below) come from numerically solving the system of ODEs describing the change in time of the concentrations of all protein, lipid, and bound species (Fig S1) via Mathematica (Equations listed in SI Text). The initial conditions had all proteins and lipids unbound and all proteins in solution.

For all simulations, our default was k_off_ rates of 1s^−1^. Then k_on_^3D^ was defined via Eq. 10. Exceptions were for proteins with known rates, and for the few simulations where to prevent k_on_^3D^ from exceeding the diffusion-limited value of 4πsD_tot_, we used k_off_=4πsD_tot_/K_a_^PP^. Although the ODEs do not use diffusion constants, we did need them to define k_a_^3D^ (Eq. 9), then k_a_^2D^ (Eq. 13), then k_on_^2D^ (Eq. 11). We used D3D=50μm2/s and D2D=0.5μm2/s for each specie, both reasonable estimates for diffusion in solution and lipid diffusion(12). For equilibrium measurements (Fig 2 and Fig 3a,b) we also simply defined k_on_^2D^=k_on_^3D^/(2σ). To calculate the percentage of proteins in complex, we used 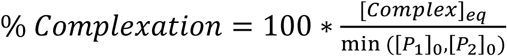

**C2. *Simulations with scaffold proteins*** For the scaffold-mediated system (Fig S7), the addition of the scaffold protein (SP) with two binding sites, one for each peripheral membrane protein, meant a total of 14 species could be formed, producing a larger system of ODEs to solve. The ODEs were solved with Mathematica or VCell. Both K_a_^eff^ and K_a_^PP^ were extracted from simulations for all systems (SI Text, Table S2), with and without membrane present, respectively. This allowed us to measure the enhancement in binding due to localization, just as for the pairs, even though here K_a_ is not a true equilibrium constant for complex formation. Detailed equations in SI Text.

**C3. *Rule-based stochastic simulations of higher order oligomers and clathrin lattice formation*** To study not only dimerization or binding mediated by a single scaffold protein, but binding of components into chain-forming oligomers or clathrin lattices, we performed Gillespie simulations(42) written in our lab using a rule-based implementation. Rule-based implementations(43) allow one to track formation of large multi-protein complexes including dimers, trimers, n-order oligomers, etc., without having to enumerate all possible complexes in advance, which is a huge challenge to encode in a system of ODEs. These simulations lack spatial or structural detail, so although we can track complexes formed, we cannot visualize assemblies or structural features. To study BAR domain proteins forming oligomers, the BAR proteins each contained their dimer forming interaction sites as well as an additional non-competing site, allowing oligomeric filaments to form. Because oligomerization was not observed in solution even at ∼100μM concentrations (11), we assume weak oligomer contacts of 500μM, which will produce <5% of proteins in higher order complexes in solution. We calculate %oligomerization as the number of bound oligomer sites on the partner at lower concentration, relative to its total concentration. Full simulation conditions in Dataset S4.

To study clathrin polymerization, each trimer leg (one clathrin molecule has three trimer legs) was able to bind to any trimer leg of another clathrin molecule, and these interactions did not compete with adaptor binding. The ability of clathrin to interact with other trimers was assumed to be independent of its interactions with the adaptor AP-2 and no cooperative binding of clathrin was included, to minimize the number of adjustable parameters and consider the simplest model of cage formation (Table S2, Fig S8). Clathrin polymerization was simulated for the *in vitro* experimental conditions reported in Kelly et al (1). We extracted a V/A ratio of 9.46μm and a lipid concentration of 54,668 μm-2 from the study.

**C4. *Spatially resolved reaction-diffusion simulations*** Single particle reaction-diffusion (RD) simulations were used to measure time-scales of assembly formation (Fig 3e, Fig S4) in a way that explicitly captured the spatial distribution of proteins and lipids and the diffusion of species to contact. We used the Free Propagator Reweighting (FPR) algorithm, an efficient and highly accurate method for studying reactions between diffusing species at spatial and single molecule resolution both in solution (19) and on the membrane (20). All lipids are initialized in the membrane plane, which is the bottom plane of the simulation box, distributed randomly. Each protein is a sphere, and binding to a lipid (also a sphere) does not prevent binding to the protein partner, and vice versa. The simulation box has periodic boundaries in the x and y dimensions, and the z dimensions are both reflective, with the lower z plane containing the reactive lipids. The equilibrium properties of the RD simulations agreed with the ODE simulations, because of the conserved equilibrium constants (Fig S4). The time-dependent properties of the RD simulations did not differ significantly from the ODEs (Fig 3e, Fig S4) due firstly because we took care in assigning corresponding macroscopic and microscopic rate constants above. Secondly, the spatial dimensions of the RD systems we simulated were small enough (box of 0.47x0.47x0.76μm) that diffusion to reach the membrane did not slow down equilibration. For box sizes with larger distances to reach the membrane, however, the RD equilibration time slows relative to the ODEs due to this spatial effect.

The FPR code for performing these RD simulations is available for download from github.com/mjohn218.

### D Collecting biochemical data, *in vivo* geometry, and concentrations

In Table 1 we list all the human and yeast proteins for which we were able to collect sufficient biochemical data on lipid and protein interactions. The 20 lipid-binding yeast proteins were retained from a larger list of 139 peripheral membrane proteins (PMP) identified from the Uniprot database as having lipid binding activity in yeast (Dataset S1). Between this set of 139 PMPs, we found 396 interactions via BioGRID, however, only 17 pairs (Table S1) involved partners with known K_a_^PM’^s. The 15 human proteins studied are all involved in CME and their biochemical data (Table 1, Table S1, Datasets S3-S4) was collected via extensive literature curation. To study scaffold-mediated interactions (Table S2), we identified all possible interactions that involved a non-membrane binding protein that could simultaneously and non-competitively bind to two of our PMPs. For the yeast proteins, these interactions could be identified from the manually curated interface interaction network for CME proteins (44). There was a relatively small number of examples where a single scaffold protein was capable of bridging two PMPs (Table S2). These interactions in humans/yeast involved clathrin/clathrin, eps15/ede1, or itsn1/sla1.

In Table S3 we collected volume and surface areas for cells and organelles with justifications provided. Because the cytoplasmic volume typically constitutes 50-60% of the total cell volume in mammalian cells, our V/A ratios set the solution volume as 60% of the total cell volume for all cell types. Lipid concentrations are collected in Table S4. The concentrations of specific lipids on specific membranes have only been quantified in a few cases, such as PI(4,5)P_2_ having an average concentration of 2.5x104μm-2 on the plasma membrane in mouse fibroblasts (15). We used this concentration as a gold standard, due to its relative consistency across measurements (13, 15), and other phosphoinositide concentrations were quantified relative to this one. We curated literature to collect the necessary copy numbers of each lipid in the cell, and their distributions across organelles. Lastly, protein concentrations were defined from copy numbers measured in yeast (45) and human cells (46) (Table 1).

## ACKNOWLEDGEMENTS

Research reported in this publication was supported by the National Institute of General Medical Sciences of the NIH under award R00GM098371 to MEJ. The content is solely the responsibility of the authors and does not necessarily represent the official views of the NIH. The research used the NSF XSEDE computational resources of Stampede and Comet, the Homewood high-performance compute cluster at JHU and the Maryland Advanced Research Computing Center clusters. We thank Doug Barrick, Attila Szabo, and Bill Eaton for helpful comments on the manuscript.

